# Sulbactam-durlobactam susceptibility among cefiderocol heteroresistant *Acinetobacter*

**DOI:** 10.1101/2025.03.05.641507

**Authors:** Bikash Bogati, Tugba Ozturk, Sarah W. Satola, David S. Weiss

## Abstract

The ATTACK clinical trial for treatment of carbapenem-resistant *Acinetobacter baumannii-calcoaceticus* complex (CRAB) isolates determined treatment with sulbactam-durlobactam to be efficacious and safe. However, other newly introduced β-lactam antibiotics, including the novel cephalosporin cefiderocol, have been compromised upon clinical introduction by a type of antibiotic resistance called heteroresistance, in which only a small subpopulation of total cells exhibit phenotypic resistance. Therefore, we sought to test for sulbactam-durlobactam heteroresistance, as well as whether sulbactam-durlobactam was effective against cefiderocol heteroresistant CRAB isolates. We did not observe heteroresistance (or conventional resistance) to sulbactam-durlobactam among the 107 carbapenem-resistant *Acinetobacter* isolates tested, consistent with the efficacy of this new antibiotic in the ATTACK trial. Further, sulbactam-durlobactam was active against cefiderocol heteroresistant CRAB, highlighting that this antibiotic may be prioritized in relation to cefiderocol in treating CRAB infections.

---

We read with great interest the manuscript by Kaye and colleagues on the ATTACK trial demonstrating the efficacy and safety of sulbactam-durlobactam in treating serious infections caused by carbapenem-resistant *Acinetobacter baumannii-calcoaceticus* complex (ABC) ^1^. A 16-fold (64 mg/mL to 4 mg/mL) average reduction in MIC_90_ of sulbactam was observed after addition of durlobactam in *in vitro* susceptibility testing of 175 baseline ABC isolates ^2^. However, standard antimicrobial susceptibility testing cannot detect heteroresistance - a form of resistance in which a minority resistant subpopulation of cells co-exists with a majority susceptible population (appendix p 3) ^3^. We previously observed a correlation between the rate of treatment failure to another recently approved beta-lactam, cefiderocol, and heteroresistance ^4^. We therefore set out to test if we would 1) identify heteroresistance to sulbactam-durlobactam and/or 2) if this novel drug could kill cefiderocol heteroresistant, carbapenem-resistant ABC.

We performed population analysis profile (PAP; appendix p 3), the gold standard test to detect heteroresistance, for sulbactam-durlobactam on 107 carbapenem-resistant *Acinetobacter baumannii* (CRAB) isolates collected between 2012 and 2015 from the CDC-funded Georgia Emerging Infections Program multi-site gram-negative surveillance initiative (MuGSI). Sixty-three of these isolates were previously shown to exhibit heteroresistance to cefiderocol ^4^. We did not observe heteroresistance (or conventional resistance) to sulbactam-durlobactam among the 107 CRAB isolates (Table). These data indicate that sulbactam-durlobactam overcomes cefiderocol heteroresistance among CRAB isolates. While treatment outcomes are dependent on multiple factors, these data align with the increased clinical activity of sulbactam-durlobactam against CRAB in the ATTACK trial as compared to that of cefiderocol in the CREDIBLE trial (all-cause mortality for cases with *Acinetobacter* infection: 19% vs 49% respectively) ^1,5^. Consistent with these data, cases of successful sulbactam-durlobactam treatment against cefiderocol resistant CRAB have been reported ^6^.

Of note, imipenem was included in both treatment arms in the ATTACK trial. An open question is whether imipenem was required to support the activity of sulbactam-durlobactam against CRAB. Our *in vitro* data demonstrated that all the isolates we tested were susceptible to sulbactam-durlobactam in the absence of imipenem, and thus that imipenem was not required for the antibacterial activity of sulbactam-durlobactam (appendix p 5).

Taken together, these data demonstrate the absence of sulbactam-durlobactam heteroresistance among our clinical isolate collection of CRAB, including those previously demonstrated to exhibit heteroresistance to cefiderocol. While these data do not imply that sulbactam-durlobactam heteroresistance does not or will not exist, nor that all cefiderocol heteroresistant CRAB isolates will necessarily be susceptible to sulbactam-durlobactam, the data are consistent with the enhanced clinical activity of sulbactam-durlobactam.

**Table.**
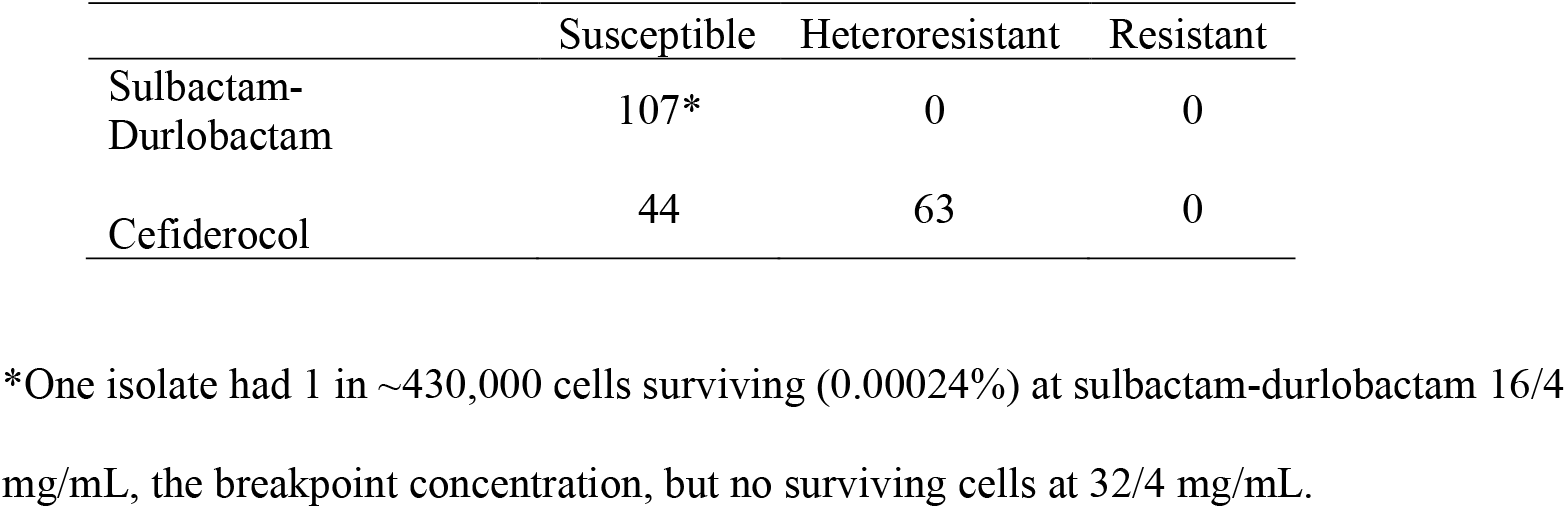
Susceptibility profile of 107 *Acinetobacter* clinical isolates based on population analysis profile.

|  | Susceptible | Heteroresistant | Resistant |
| --- | --- | --- | --- |
| Sulbactam-Durlobactam | 107* | 0 | 0 |
| Cefiderocol | 44 | 63 | 0 |

\*One isolate had 1 in ~430,000 cells surviving (0.00024%) at sulbactam-durlobactam 16/4 mg/mL, the breakpoint concentration, but no surviving cells at 32/4 mg/mL.

## Acknowledgements

This research was supported by NIH grant AI158080 and Department of Veteran’s Affairs grant BX005985 to DSW. DSW was also funded by a Burroughs Wellcome Fund Investigator in the Pathogenesis of Infectious Disease award.

## Author contributions

BB and DSW conceived of the study. BB and TO performed all experiments. BB, TO, SWS, and DSW conferred on study design and data analysis. BB wrote the manuscript with assistance from TO, SWS, and DSW.

## METHODS

### Isolate information

Carbapenem-resistant *Acinetobacter baumannii* (CRAB, from 2012-2015) were collected in Georgia, USA by the CDC-funded Georgia Emerging Infections Program.

### Population analysis profile

Population analysis profile (PAP) was performed as described previously ^1^. The CLSI breakpoint for CRAB for sulbactam-durlobactam is 16-4 *μ*g/mL and that for imipenem is 8 *μ*g/mL ^2^. CRAB isolates were grown overnight from a single colony isolated from frozen stock in 1.5 mL Mueller-Hinton broth at 37°C with shaking at 200 rpm. The culture was then serially diluted and plated on Mueller-Hinton agar containing 0, 2/4, 4/4, 8/4, 16/4, 32/4 mg/mL of sulbactam (Thermo Fisher Scientific)-durlobactam (MedChemExpress), 0, 2, 4, 8, 16 mg/mL of imipenem (Biosynth International), or 0, 2/4/4, 4/8/4, 8/16/4, 16/32/4 mg/mL of imipenem-sulbactam-durlobactam. Colonies were enumerated after 24-48 h of growth at 37°C. An isolate was classified resistant if the number of colonies that grew at the breakpoint concentration was at least 50% of those that grew on plates without antibiotic. If an isolate was not resistant, it was classified heteroresistant if the number of colonies that grew at 1 or 2 times the relevant breakpoint was at least 0.0001% (1 in 10^6^) of those that grew on plates without antibiotic. If an isolate was neither classified resistant nor heteroresistant, it was classified susceptible. PAP results for cefiderocol were reported previously ^3^.

## Supplemental Figure

**Supplemental Figure.**
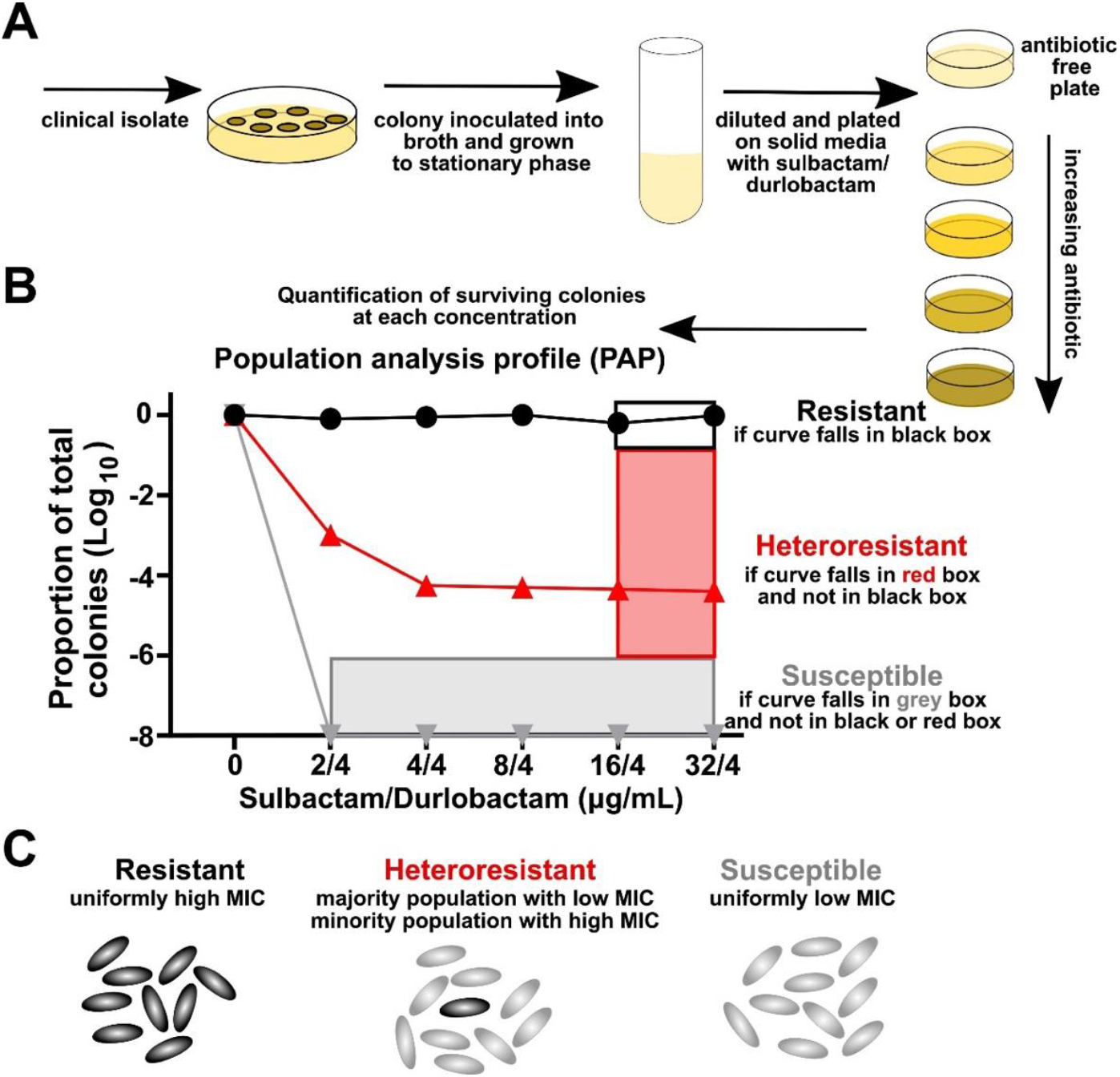
Overview of population analysis profile and heteroresistance. (A) An *Acinetobacter* isolate is isolated and grown in liquid medium, and then serially diluted and grown on increasing concentrations of sulbactam-durlobactam. (B) Surviving colonies are enumerated and the isolate is classified as resistant if at least 50% of the total colonies grow at 1x or 2x the breakpoint concentration. An isolate is considered susceptible if less than -6 logs (0.0001%) grow at 1x or 2x the breakpoint concentration. An isolate is considered heteroresistant if there is less than 50% survival at 1x breakpoint and greater than 0.0001% at 2x breakpoint concentration. (C) A schematic depiction of the cells grown from a single colony of a resistant, heteroresistant, or susceptible isolate. Resistant cells shown in black and susceptible cells shown in grey.

**Supplemental Table.**
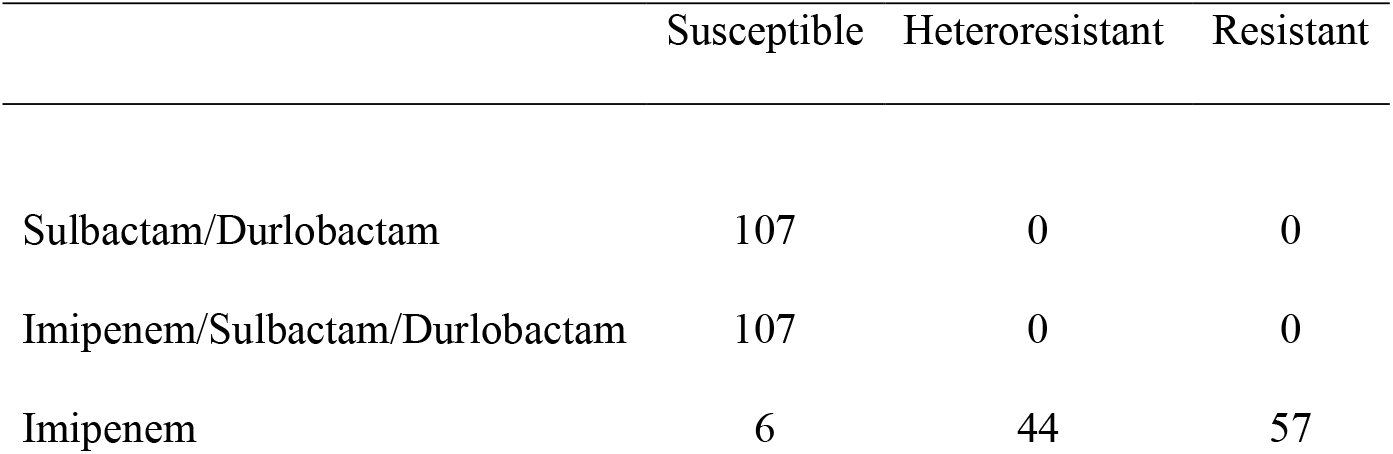
Susceptibility profile of 107 *Acinetobacter* strains based on population analysis profile.

|  | Susceptible | Heteroresistant | Resistant |
| --- | --- | --- | --- |
| Sulbactam/Durlobactam | 107 | 0 | 0 |
| Imipenem/Sulbactam/Durlobactam | 107 | 0 | 0 |
| Imipenem | 6 | 44 | 57 |

